# Succession and Shifting Identities in Freshwater, Built Environment Biofilm Communities

**DOI:** 10.64898/2026.06.23.734043

**Authors:** Todd Testerman, Stacy King, Timothy J. Welch, Gregory D. Wiens, Joerg Graf

**Affiliations:** University of Connecticut, Department of Molecular and Cell Biology, Storrs, Connecticut, USA; Riverence Provisions, Buhl, Idaho, USA; National Center for Cool and Cold Water Aquaculture, Agricultural Research Service/U.S. Department of Agriculture, Kearneysville, West Virginia, USA

**Keywords:** biofilm, succession, aquaculture, hatchery, 16S rRNA, amplicon sequencing, rainbow trout, freshwater, built environment

## Abstract

Biofilms on aquaculture infrastructure harbor diverse microbial communities that may influence water quality and fish health, yet the temporal dynamics of these communities remain poorly characterized. Here, we used 16S rRNA gene amplicon sequencing to profile biofilm communities on concrete raceway surfaces across an 80-day rainbow trout (*Oncorhynchus mykiss*) indoor hatch-house production period. One hundred twenty-three wall swab samples from 19 raceways at six time points (9, 23, 38, 53, 65, and 80 days) were analyzed after stringent quality control. Beta diversity analyses revealed that biofilm communities at each time point were significantly distinct (PERMANOVA, p < 0.001 for all pairwise comparisons), with early communities exhibiting greater variability than late-stage biofilms. Total bacterial load increased approximately 2.5-fold from early to late stages (qPCR, p < 0.001). Differential abundance testing (ANCOM-BC) identified 57 differentially abundant genera between early-and late-stage biofilms, and random forest classification distinguished early from late communities with over 93% test accuracy. A clear successional trajectory emerged: early biofilms were dominated by pioneer taxa including *Pseudomonas, Caulobacter, and Flavobacterium*; mid-succession communities featured predatory *Bdellovibrio* and the methylotroph *Methylotenera*; and mature biofilms were enriched in saprophytic Saprospiraceae and *Haliscomenobacter*, polysaccharide-degrading Verrucomicrobiaceae, and cooperative predatory myxobacteria. *Flavobacterium columnare*, a pathogen of concern in aquaculture, was detected at low levels throughout the production period. These results demonstrate predictable ecological succession in freshwater built environment biofilms and provide a foundation for understanding the role of surface-associated microbial communities in hatchery management.

## Introduction

The colonization of a surface by microbes begins the process of community assembly and biofilm formation. This initial attachment phase sees microbial members from the seeding community adhere, proliferate, and manipulate the local microenvironment through the production and secretion of extracellular polymeric substances (EPS), nucleic acids, primary and secondary metabolites, and waste products (1–4). These so-called founder species play an important role in influencing the maturing biofilm through niche modification and competitive mechanisms (5, 6). As the biofilm continues to develop, new constituents enter the community, further increasing diversity while often displacing or reducing the abundance of founder species (7). Eventually space is exhausted, and a mature biofilm begins to shed constituents leading to dispersal of member cells to other environments (2, 8).

This process has been studied *in vitro* through the use of confocal microscopy and flow cells which allow for real time monitoring of biofilm development (9, 10). Additionally, *in vivo* profiling of microbial community assembly on surfaces in a variety of environments including marine (11–13), freshwater (14, 15), sewage systems (16), and others (17, 18) have displayed the interesting variability and, often, unpredictability of microbial community assemblages. These biofilm communities present serious problems and provide interesting solutions as they relate to the processes of biofouling and bioremediation (19, 20). Biofilms also act as reservoirs for pathogenic microorganisms (21–24) and have been shown to facilitate the spread of antimicrobial resistance (AMR) (25–27), highlighting the importance of aquatic biofilms in human healthcare and thus for One Health (28, 29).

In aquaculture systems, biofilms forming on infrastructure surfaces have received growing attention for their roles in water quality maintenance and disease ecology. In recirculating aquaculture systems (RAS), biofilter biofilms are essential for nitrification and ammonia removal, and their microbial community composition has been characterized across a range of production systems (30, 31). Beyond biofilters, biofilms colonizing rearing infrastructure (tank walls, screens, pipes, and other surfaces in direct contact with fish) harbor diverse bacterial communities that can include opportunistic fish pathogens (32, 33). An initial multi-year survey of microbial communities at the facility examined in the present study characterized biofilm and planktonic assemblages across surface types and environmental conditions, and demonstrated that the fish pathogen *Flavobacterium columnare* resides within abiotic surface biofilms and is seeded from the natural water source (34). However, most studies of aquaculture-associated biofilms have characterized communities at single time points or across broad temporal scales. The fine-scale successional dynamics of biofilm community assembly within a single production period (the period beginning when water and fish fry are added to a raceway and ending once the raceway is drained and the fish are moved outdoors), from initial surface colonization through biofilm maturation, remain largely unexplored. Understanding these dynamics is important because biofilm management decisions such as cleaning frequency and chemical treatment regimens are currently made without knowledge of how biofilm communities develop over time or at what stage they might become problematic or potentially beneficial.

In the present study, we sought to characterize the successional dynamics of biofilm communities on built surfaces within a freshwater aquaculture environment. Wall swabs were collected from 19 individual rainbow trout (*Oncorhynchus mykiss*) raceways within an indoor hatchery facility, the same facility described in an initial characterization of hatchery surface communities (34). Biofilms on these sampled surfaces ranged in maturity from 9 to 80 days, spanning a complete indoor early-life stage production period, that occurs prior to the transfer to outdoor raceways for grow out. Through the use of 16S rRNA gene sequencing, we probed the shifting compositional structure of these surface biofilms over time and examined the potential for biofilm community development to impact the abundance of a known fish pathogen, *Flavobacterium columnare*.

## Materials and Methods

### Site Description

Samples were collected from raceways located in a rainbow trout hatchery building in Idaho, USA in 2021. The hatchery building is an entirely enclosed structure with minimal natural light and has 20 epoxy-coated concrete raceways supplied with water from an upstream spring (Supplementary Figure 1). Extensive sampling of 19 separate raceways at a single time point was conducted in August. The water temperature at all sampling times was 14-15°C. Surfaces within hatchery raceways are regularly scrubbed with a coarse brush with the frequency increased with fish density as per standard fish husbandry practice. The precise timing of bottom scrubbing in relation to samplings was not recorded. The water level within each hatchery raceway was increased periodically as the fish continued to grow. Following the 80-day indoor production period, each hatchery raceway was drained, pressure washed, soaked in a bleach solution (20%), and then dried. The available water quality parameters of the influent were: 0.017 mg/L total phosphorus and <2 mg/L total dissolved solids.

### Sampling of Surfaces

Three surface types were sampled within each raceway: the side walls, the tailscreens, and the baffles. The side walls were the primary focus of this study because they represent the largest continuous submerged surface area within a raceway, are of uniform material (epoxy-coated concrete), and are present in a directly comparable form across all raceways, providing a standardized substrate for comparing biofilm succession between raceways. The tailscreens and baffles, which present smaller and structurally distinct surfaces, were also sampled and are reported for comparison in the Supplementary Material (Supplementary Figures 2 and 3). The raceway floor was not sampled, because settleable solids and feces accumulate on the bottom; swabbing there would have captured trapped particulate material and settled planktonic cells rather than the surface-attached biofilm that was the focus of this study. Hatchery surfaces were swabbed as previously described (34), with each surface swabbed until visible biofilm discoloration was removed rather than over a fixed surface area. Within each raceway, the wall was swabbed at two positions along its length, one within the first third and one within the final third of the raceway. At each position, a vertical series of swabs was collected at successive depths, from the air-water interface to the base of the wall, with sampling depth recorded in swab-length increments below the waterline. All swabs were placed on ice following sampling and then frozen at -20°C for storage and shipped on dry ice.

### DNA Extraction, PCR Amplification, Library Preparation, and Sequencing

DNA was isolated from surface swab samples and the V4 hypervariable region of the 16S rRNA gene was amplified as previously described (34). The resulting completed PCR reactions were prepared for sequencing as previously described (35).

### Bioinformatic Processing

Demultiplexing of 16S V4 rRNA gene sequence data was done through Basespace (basespace.illumina.com). Paired-end data was then downloaded and imported into QIIME 2 (2024.10) (36) for filtering, denoising, dereplication, and amplicon sequence variant (ASV) calling via the DADA2 denoise-paired plugin (37). Sequences were classified taxonomically using the feature-classifier (38) and taxa (https://github.com/qiime2/q2-taxa) plugins with a Naïve Bayes classifier based on the Silva 13899% OTU database (https://www.arb-silva.de/documentation/release-138/) (39) trimmed to the region bound by the 515F/806R primer pair used as a reference. Default QIIME 2 processing parameters were used in this workflow except where otherwise noted in the command logs.

The feature table, taxonomy table, and rooted phylogenetic tree were imported into RStudio (2024.12.0+467) (40) from QIIME 2 using the qiime2R package (41). Phyloseq (v1.50.0) (42), ggplot2 (v3.5.1), microViz (v0.12.6) (43), and ANCOM-BC (analysis of composition of microbiomes with bias correction; v2.8.0) (44) were employed for alpha and beta diversity calculation and barplot, boxplot, and ordination figure generation. Statistical testing was performed using the vegan (v2.6.8) (45) and rstatix (v0.7.2) packages.

### Quality Control

Controls were processed for every DNA extraction kit lot to assess extraction kit contaminants and contamination occurring during DNA extraction. PCR controls were run for every PCR plate using molecular grade water instead of DNA to assess PCR reagent contaminants as well as contamination occurring during PCR setup. Mock community DNA controls (ZymoBIOMICS, Irvine, CA, USA; cat. # D6306) were also amplified during each PCR and used to assess PCR amplification biases.

During library preparation, a QIAxcel was used to verify successful PCR reactions as well as expected amplicon size. A qPCR assay was used to quantify the total 16S rRNA gene copy number in each sample using a previously validated assay (46) to verify that total bacterial load was great enough to negate the effects of well-to-well contamination. To improve the robustness of our dataset, we excluded samples with either 8,000 or fewer 16S copies (as determined by 16S qPCR) or a sequencing read depth of 8,000 reads or less (289 samples retained/528 samples sequenced, 54.7% retained). Between these two quality control metrics, no negative controls were retained. Quality filtering and chimeric sequence removal was performed using QIIME2 with default parameters.

Total bacterial load in each sample was estimated by quantitative PCR (qPCR) targeting the 16S rRNA gene using eubacterial primers 27F and 519R, as previously described (34). Reactions contained 2x SsoFast EvaGreen Master Mix (Bio-Rad), primers at 300 nM final concentration, 5 µL of template DNA, and PCR-grade water to a final volume of 20 µL. Thermocycling was performed on a CFX96 Touch real-time thermocycler (Bio-Rad) with the following conditions: 98°C for 3 min, followed by 40 cycles of 98°C for 10 s and 55°C for 30 s, with a plate read step after each cycle. Quantification cycle (Cq) values were determined using baseline-subtracted curve fitting. Samples with fewer than 8,000 16S rRNA gene copies were excluded to minimize the influence of well-to-well contamination inherent to plate-based PCR setups.

### Statistical Analysis

Diversity metrics were calculated using data rarefied to 8,000 reads. Permutational Analysis of Variance (PERMANOVA) (47) testing was used to determine statistical significance of group differences for relevant beta diversity metrics. To assess whether sampling depth structured wall biofilm communities, PERMANOVA was additionally performed with sampling depth as a factor, both alone and with biofilm age or raceway identity included as a covariate, and homogeneity of multivariate dispersion among depths was evaluated with beta dispersion testing. ANCOM-BC (44) was used for differential abundance testing with default parameters. The structural zero flag was set to “True” for comparisons only where both groups had 30 or more samples each, as recommended. A corrected p-value of 0.05 was required for significance to be achieved. Shapiro-Wilk normality testing was performed on each alpha diversity metric on a per group basis. Normally distributed metrics were compared using pairwise t-tests and non-normally distributed metrics were compared using Wilcoxon tests. P-values were corrected due to multiple comparisons using the Holm method for these aforementioned tests. Total bacterial load was estimated from 16S rRNA gene copy numbers determined by qPCR. Differences in total bacterial load across time points were assessed using Kruskal-Wallis tests with pairwise Wilcoxon comparisons (Holm-corrected), and the relationship between biofilm age and total bacterial load was evaluated using Spearman rank correlation. Random forest classification was performed using the randomForest package (v4.7-1.2) with the rsample package (v1.2.1) used for train/test data splitting. 16S rRNA qPCR data were used to assess trends in total bacterial load but were not used to estimate taxon-specific absolute abundances, as variability in per-taxon 16S gene copy number and swab-to-swab sampling efficiency preclude reliable conversion.

## Results

### Surface Microbial Communities are Temporally Distinct

One hundred twenty-three wall swabs from 19 hatchery raceways at 6 distinct time points remained after quality control and were analyzed. Figure 1 shows the relative abundances of reads for each sample grouped by time point at the class and genus levels. At the class level, the Bacteroidia (median, 33.5%), Alphaproteobacteria (26.0%), and Gammaproteobacteria (22.0%) predominate throughout the production period with the Verrucomicrobiae (4.7%) increasing at later time points. At the genus level, the communities appear distinct upon visual inspection with the day 9 communities appearing to be very different than the day 80 communities. In the earliest biofilms, members of the Comamonadaceae family and genera *Flavobacterium* and *Rheinheimera* comprise approximately 50% of the communities. As time passes, community diversity increases and the relative abundance of members of the Saprospiraceae family increases as well. Community profiles for tailscreen and baffle surfaces showed similar taxonomic compositions (Supplementary Figures 2 and 3).

Beta diversity metrics of these groups were compared to determine if the microbial communities differed significantly between time points. Figure 2 displays the Bray-Curtis (A), Jaccard (B), and weighted (C) and unweighted UniFrac (D) NMDS ordinations comparing these samples. Distinct clustering of samples can be noted when comparing the youngest (9 days) and oldest (80 days) biofilm communities, with a clear temporal gradient visible for all metrics. A principal component analysis biplot further highlighted the top genera driving community separation across time points (Supplementary Figure 4). Subsequent PERMANOVA testing for the variable of maturity returned significant results (p.adj < 0.001) for the four metrics shown. Additionally, pairwise PERMANOVA testing with multiple testing corrections between each age group returned significant results for all comparisons (p.adj < 0.001) (Supplementary Tables 1-4). Visually, it was noted that the size of the 95% confidence ellipses varied for different maturity groups, possibly indicating differences in community variability between these groups. To assess this, beta dispersion testing was used to compare the average distances to the centroids for each group, in turn providing needed context for the significant PERMANOVA results. Significant differences were noted between some of the groups for the four metrics, with a commonly observed trend being that later stage biofilms were significantly less variable than early-stage biofilms (Supplementary Tables 5-8). Sampling depth, by contrast, explained only a small fraction of community variation (PERMANOVA R² = 0.04-0.07 across the four distance metrics; p = 0.0001), well below the variation explained by biofilm age (R² = 0.15-0.30) or raceway identity (R² = 0.30-0.52), and multivariate dispersion did not differ among depths (p = 0.70). Wall biofilm communities were therefore structured far more strongly by time than by position on the wall, supporting the use of swabs spanning the sampled depth range as replicates within each raceway.

**Figure 1:**
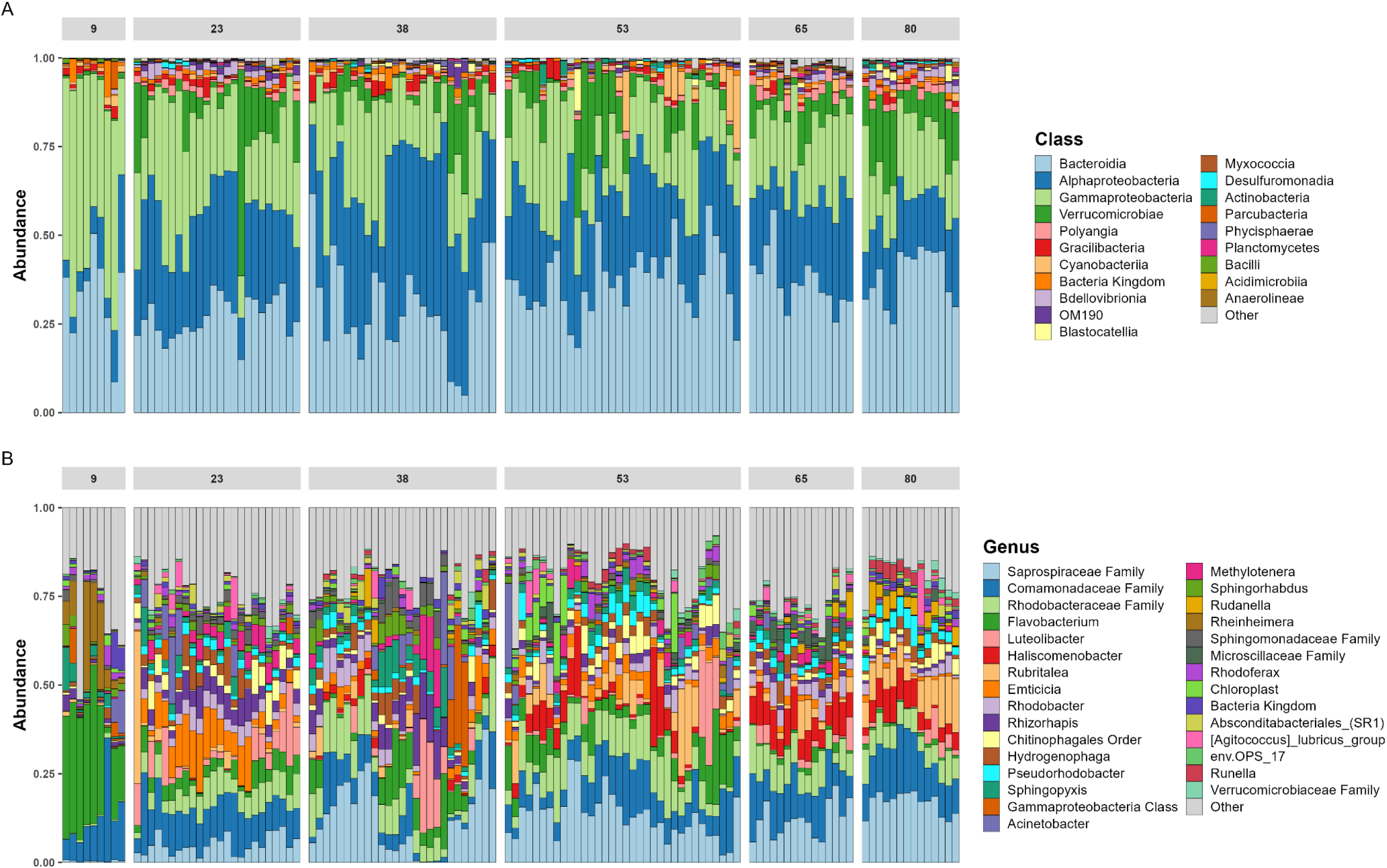
Bar Plots of Wall Biofilm Communities at the Class and Genus Level. (A) class level and (B) genus level community profiles are displayed and faceted by the number of days into the production period. Relative abundances are shown. Samples within an individual facet are grouped by their Bray-Curtis similarity. If a taxonomic rank is denoted after an entry in the legend, e.g., “Saprospiraceae Family”, this indicates that this grouping of reads could not be classified beyond the family level. This could represent multiple genera and species but cannot be discerned using only this region of the 16S rRNA gene and this database.

**Figure 2:**
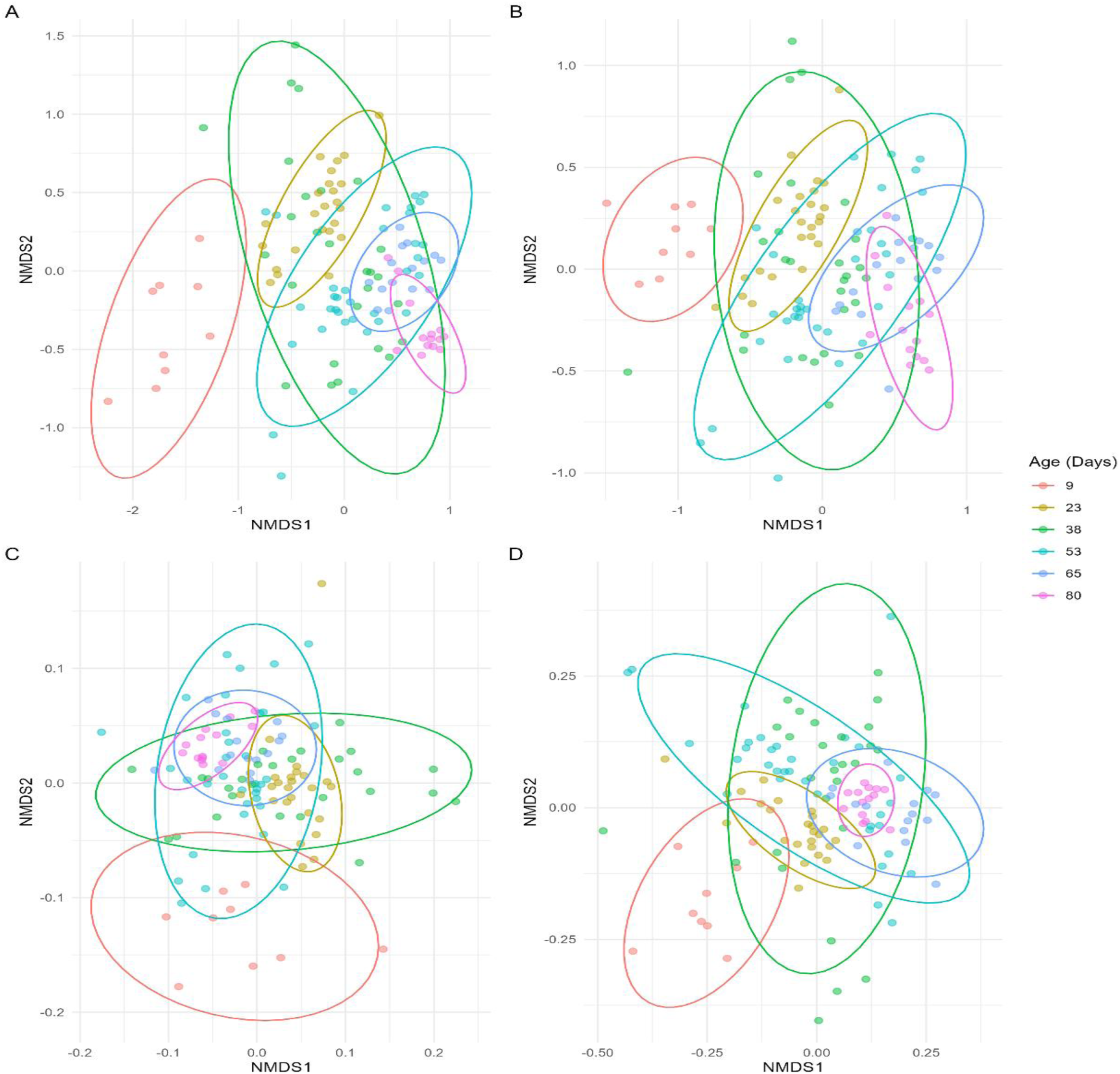
Beta Diversity by Maturity. (A) Bray, (B) Jaccard, (C) Weighted, and (D) Unweighted UniFrac NMDS ordinations are shown. Samples were rarefied to 8,000 reads before distance calculations were performed. 95% confidence ellipses are displayed for each time point. PERMANOVA results show significant differences for all metrics for maturity overall and for all pairwise testing (day 9 vs. day 80, for example).

### Richness, Evenness, and Phylogenetic Diversity Do Not Alone Explain Community Differences

With significant differences between these groups evident, we next wanted to probe potential community-level differences by comparing alpha diversity values. Figure 3 shows the observed ASVs, Shannon diversity, and Faith’s phylogenetic diversity (PD) for each maturity grouping. Most obvious is the lower alpha diversity values shown for the day 9 biofilm communities compared with all other groupings. Statistical testing comparing day 9 communities to all other communities showed significant differences for all comparisons using Shannon diversity (p.adj < 0.05) (Supplementary Table 9). For observed ASVs and Faith’s PD, only some of the comparisons yielded significant differences (Supplementary Table 9). The day 65 biofilms had consistently greater alpha diversity values than all other groups with almost all comparative testing returning significant results across the three metrics (p.adj < 0.05). Additionally, for Shannon diversity specifically, the earliest biofilms (day 9) had a significantly lower index value than all other time points (p.adj < 0.05).

**Figure 3:**
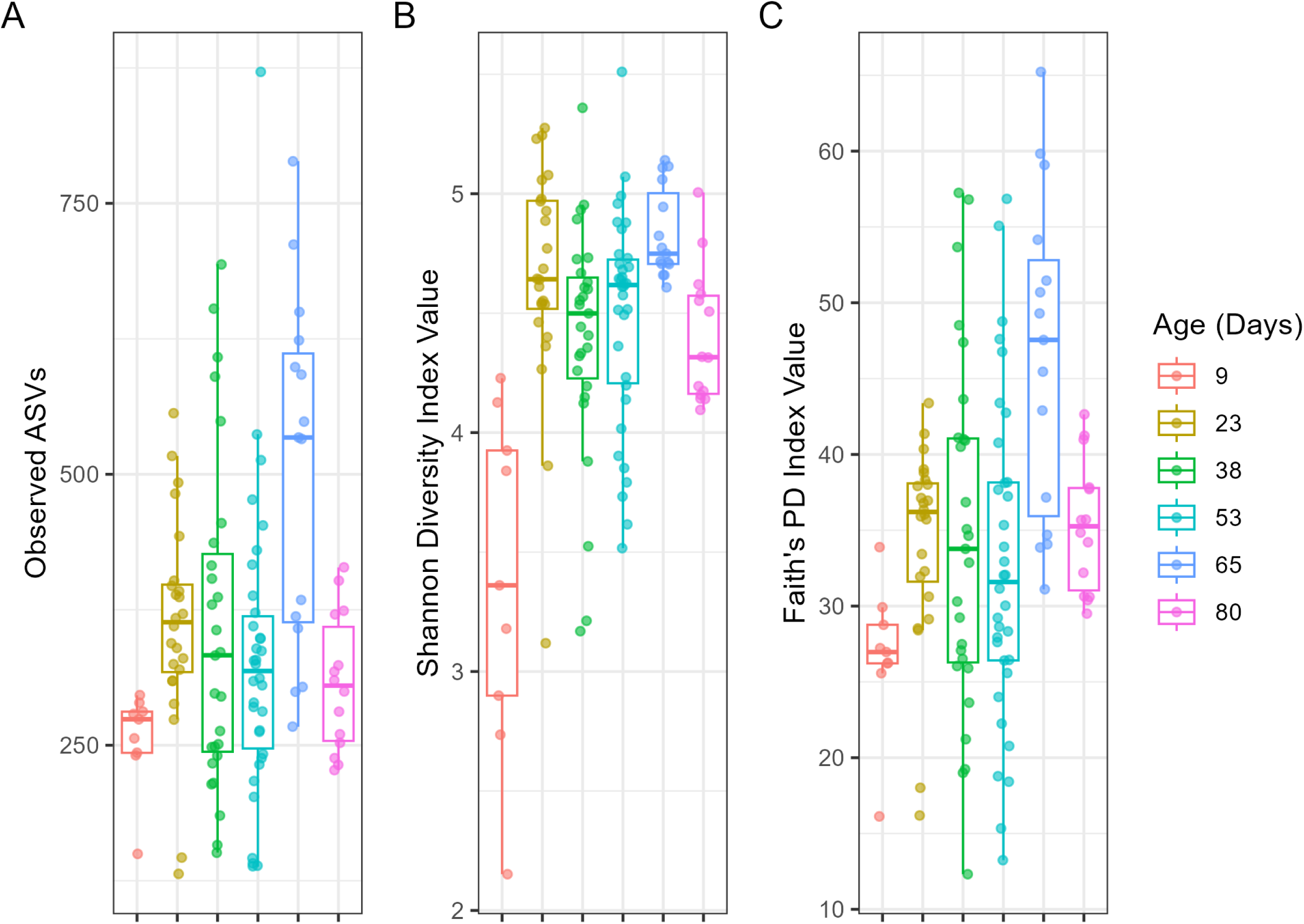
Alpha Diversity by Maturity. (A) Observed ASVs, (B) Shannon diversity, and (C) Faith’s PD are shown. Shapiro-Wilk normality testing was used to assess normality of group distributions and an appropriate follow-up test was used to compare maturity groups (T-tests or Wilcoxon tests).

To complement diversity analyses with a measure of total community size, 16S rRNA gene copy numbers from qPCR were compared across biofilm maturity as a proxy for bacterial load. Because swabbing was performed until visible discoloration rather than over a standardized surface area, and because 16S rRNA gene copy number varies among taxa, these values represent estimates of relative bacterial density rather than precise *in situ* measurements. Total bacterial load differed significantly across the six time points (Kruskal-Wallis, χ² = 37.28, p < 0.001) and was positively correlated with biofilm age (Spearman ρ = 0.50, p < 0.001). Late-stage biofilms harbored approximately 2.5-fold more bacteria than early-stage biofilms (median 633,000 vs. 251,000 16S copies; Wilcoxon, p < 0.001). Day 65 biofilms had the highest median bacterial load (1,069,000 copies), approximately five-fold greater than day 9 biofilms (213,000 copies). These data indicate that the biofilm community expands in absolute terms over the production period, and that observed shifts in relative abundance largely reflect genuine ecological turnover rather than purely compositional artifacts.

### Unique ASVs Are Present at Individual Time Points and Display Taxonomic Trends

While relative levels of different taxonomic groups clearly vary over time, we wanted to see if specific sequences were present or absent at specific time points, possibly indicating arrival or departure of individual strains of microbes. To parse these patterns, upset plots were used to track individual ASVs shared between different time points (Figure 4). The largest group of shared ASVs were between all time points between 23 and 80 days. Over 300 unique ASVs were not found in samples from day 9 biofilms and many of these ASVs belong to some of the most common taxonomic groups comprising the biofilm community (Saprospiraceae, Comamonadaceae, *Flavobacterium*). This indicates that despite many of these taxonomic groups being present at all time points, different ASVs within these taxonomic groups appear and disappear throughout biofilm development. This is exemplified when noting that specific time points have a large number of unique ASVs that are only present at that specific time point (day 65 – 230 ASVs, day 23 – 170, day 38 – 170, day 53 – 125, day 9 – 70, day 80 – 25).

**Figure 4:**
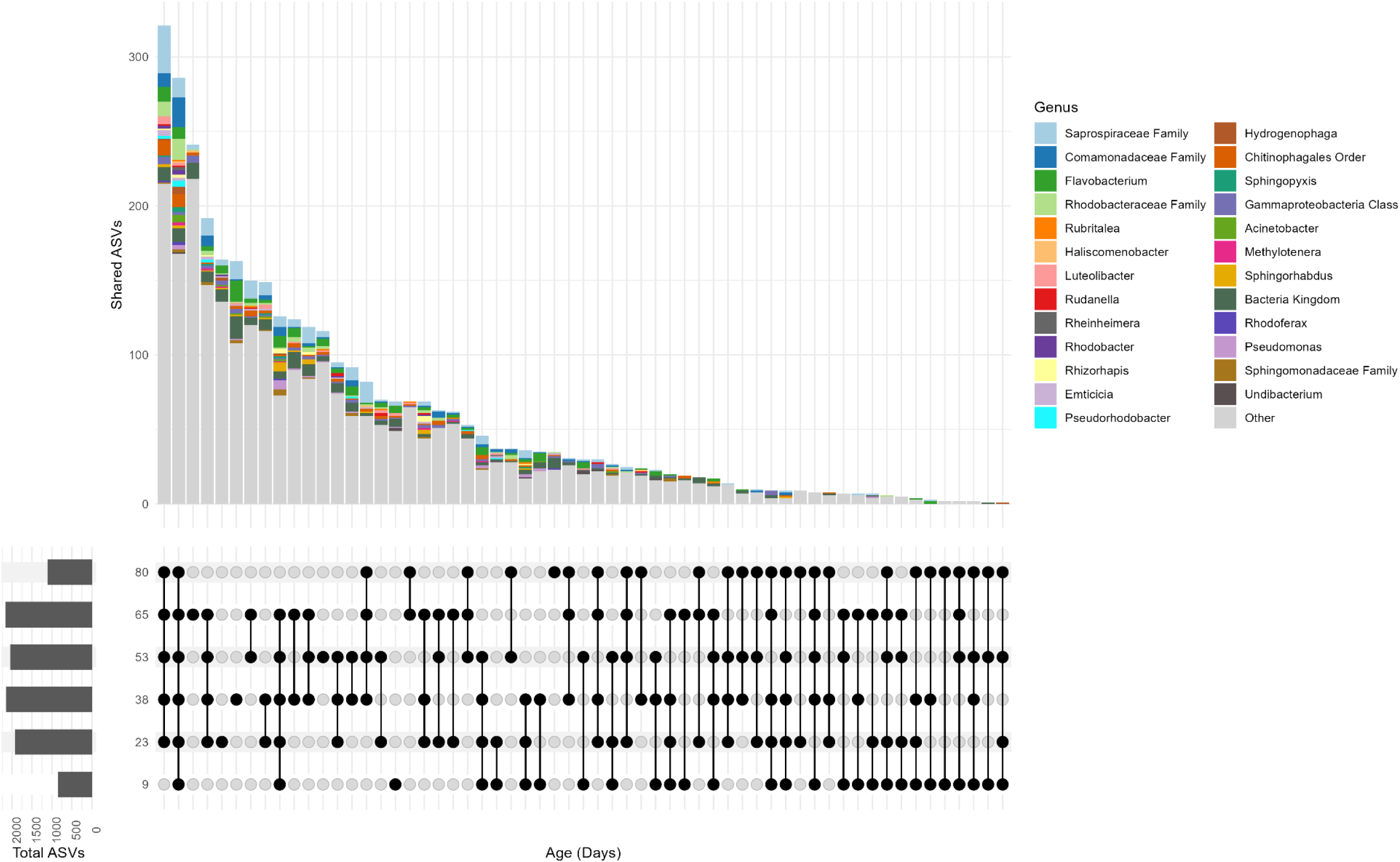
UpSet Plots Showing Shared and Unique ASVs. Values shown in the upper bar plots are the number of ASVs shared between the maturity groups corresponding to what is listed directly below the bar plot. For example, the first entry displays ASVs shared between every time point **except** for 9 days while the second entry indicates ASVs shared by **all** time points. The upper bar plots are colored by genus assignment of each ASV. The bar plots on the left show the total number of ASVs included in this comparison for each individual maturity group. To account for uneven sample numbers and uneven sampling depth, samples were merged by maturity group (all ASVs added together) and then rarefied to the lowest groups total read count. The large grey portions at the base of all bars represent all other taxonomic groups grouped together (“Other”).

Examining the taxonomic compositions of these shared or unique ASVs for all time points, specific enrichment patterns can be noted. For example, ASVs that were unique to one specific time point rarely ever belonged to the Comamonadaceae (dark blue) whereas many of the shared ASV groupings included a large proportion of Comamonadaceae. This might indicate that these specific ASVs show up in the early stages of biofilm formation and stay throughout the entire measured phase of biofilm development. Alternatively, unique ASVs belonging to the Saprospiraceae (light blue) appear at almost every individual time point possibly indicating that while this taxonomic family is present at high levels throughout biofilm development, the identity of the ASVs at lower taxonomic ranks of this family is quite dynamic and changes during maturation of the biofilm. A class-level UpSet plot showed concordant patterns (Supplementary Figure 5).

### Differential Abundance Testing Reveals Taxonomic Shifts Between Early and Late Biofilms

To identify specific taxa driving the compositional differences between early and late biofilm communities, differential abundance testing was performed using ANCOM-BC. Samples were grouped as early-stage (days 9, 23, and 38; n = 60) or late-stage (days 53, 65, and 80; n = 63) and compared at both the class and genus levels. Classes and genera not present in at least 10% of samples were excluded from this analysis, leaving 45 classes and 280 genera for testing.

At the class level, 13 of the 45 tested classes were identified as differentially abundant between early and late biofilms (Figure 5A). Four classes were significantly enriched in late-stage biofilms: WPS-2 (log fold change [LFC] = 1.27), Gemmatimonadetes (LFC = 0.98), Myxococcia (LFC = 0.81), and Thermoanaerobaculia (LFC = 0.70). Nine classes were significantly enriched in early-stage biofilms, with Bacilli showing the strongest enrichment (LFC = -1.60), followed by vadinHA49 (LFC = -1.34) and an unresolved Bacteria kingdom-level group (LFC = -1.11). The Gammaproteobacteria (LFC = -0.70) and Alphaproteobacteria (LFC = -0.59) were also enriched in early-stage biofilms, consistent with their higher representation in day 9 communities.

**Figure 5:**
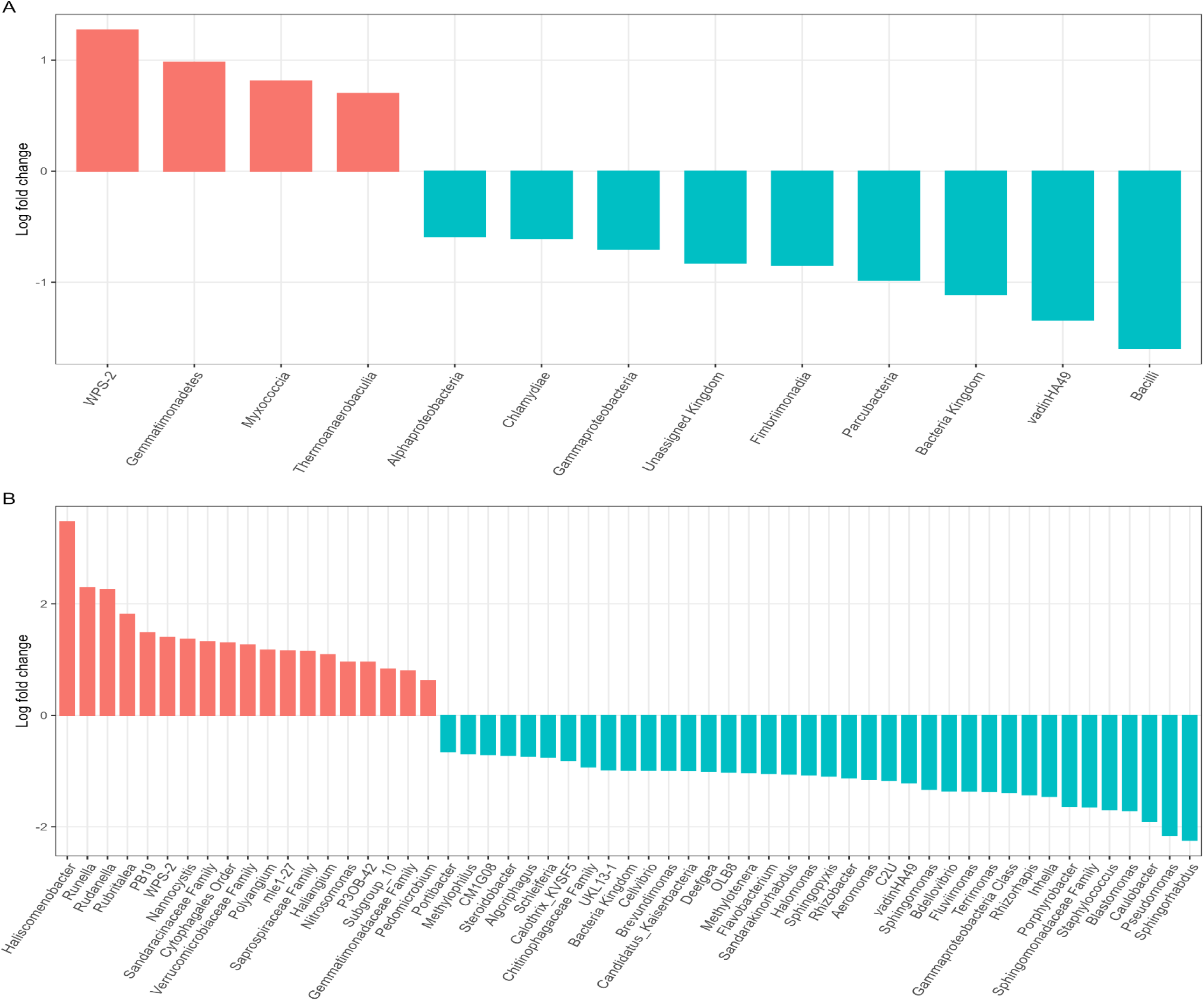
ANCOM-BC Waterfall plots comparing early and late wall swabs at class and genus levels. (A) Class and (B) genus log-fold changes are shown for taxa that were noted as differentially abundant in early (blue) or late (red) stage biofilms. Data was aggregated to the class and genus level for each respective plot. Classes and genera not appearing in 10% of the samples were excluded from this analysis.

At the genus level, 57 of the 280 tested genera were differentially abundant, with 19 enriched in late-stage and 38 enriched in early-stage biofilms (Figure 5B). *Haliscomenobacter* exhibited the largest late-stage enrichment (LFC = 3.47), followed by *Runella* (LFC = 2.29), *Rudanella* (LFC = 2.25), and *Rubritalea* (LFC = 1.81). Other notable late-enriched taxa included the predatory myxobacterium *Nannocystis* (LFC = 1.37) and related *Polyangium* (LFC = 1.17) and Haliangium (LFC = 1.09), as well as members of the Saprospiraceae (LFC = 1.15), Verrucomicrobiaceae (LFC = 1.26), and Cytophagales (LFC = 1.30) at higher taxonomic levels. The nitrifier *Nitrosomonas* was also enriched in late-stage biofilms (LFC = 0.95).

Among early-enriched genera, *Sphingorhabdus* showed the strongest enrichment (LFC = -2.24), followed by *Pseudomonas* (LFC = -2.16) and *Caulobacter* (LFC = -1.90). Additional early-enriched genera included *Inhella* (LFC = -1.45), *Rhizorhapis* (LFC = -1.42), *Porphyrobacter* (LFC = -1.63), and *Blastomonas* (LFC = - 1.71), as well as several members of the Sphingomonadaceae family (LFC = -1.65) including *Sphingomonas* (LFC = -1.33) and *Sphingopyxis* (LFC = -1.09). *Flavobacterium* (LFC = -1.04) and *Aeromonas* (LFC = -1.15) were also significantly enriched in early biofilms. The predatory bacterium *Bdellovibrio* was enriched in early communities (LFC = -1.36).

These results are largely concordant with trends observed in the beta diversity ordinations. Taxa such as *Haliscomenobacter*, Saprospiraceae, and Verrucomicrobiaceae that increase in relative abundance at later time points (Figure 6) were also identified as significantly enriched by ANCOM-BC. Similarly, early-enriched genera including *Caulobacter*, *Inhella*, and *Pseudomonas* correspond to taxa identified as important indicators of early biofilm communities by random forest (Supplementary Figure 6). The asymmetry in the number of early-enriched (38) versus late-enriched (19) genera, despite late-stage biofilms having greater alpha diversity, suggests that many of the taxa unique to late biofilms were below the 10% prevalence threshold applied here.

**Figure 6:**
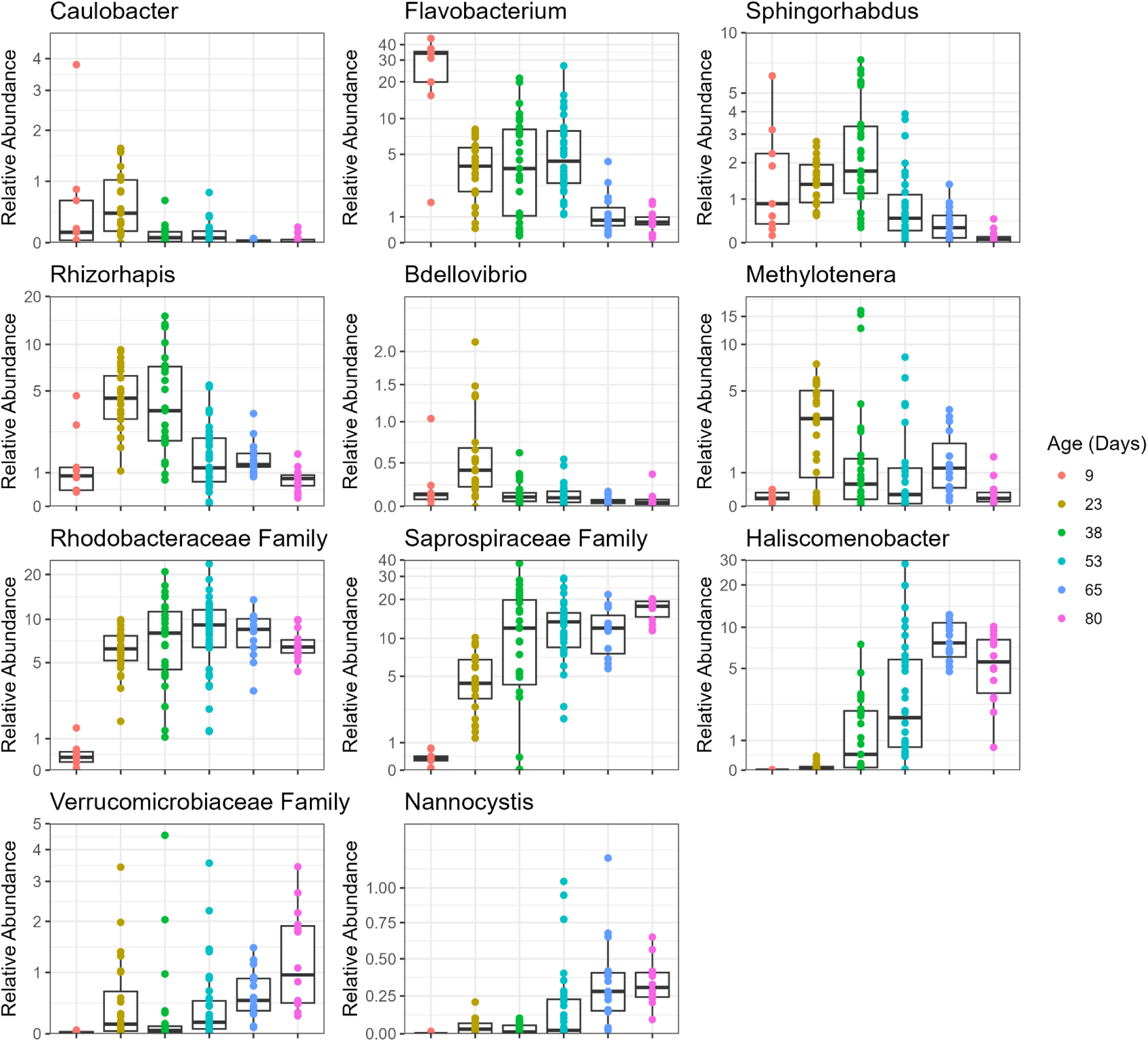
Temporal Abundance Patterns of Key Indicator Taxa. Relative abundances are shown for 11 taxa identified as important indicators of biofilm maturity by ANCOM-BC, random forest classification, or both, ordered from earliest to latest successional stage. Top row: early-stage taxa that decline with maturity (*Caulobacter*, *Flavobacterium*, *Sphingorhabdus*). Second row: mid-succession taxa peaking at day 23 (*Rhizorhapis*, *Bdellovibrio*, *Methylotenera*). Third row: persistent colonizer (Rhodobacteraceae) and late-accumulating taxa (Saprospiraceae, *Haliscomenobacter*). Bottom row: additional late-stage taxa (Verrucomicrobiaceae, *Nannocystis*). Samples are grouped by time point along the x-axis. Y-axes use pseudo-log scaling with taxon-specific breaks to accommodate differences in abundance range. If a taxonomic rank is present after the name (e.g., Saprospiraceae Family), this indicates that sequences could not be classified beyond that rank.

### Biofilm Maturity Can Be Reliably Classified Using Random Forest

After observing clear, statistically significant differences between different biofilm stages through beta diversity and differential abundance analyses, an effort to identify the important indicator taxa for various early and late-stage biofilms was undertaken. Early or late-stage samples were subset into training and test datasets. To simultaneously identify important taxa and produce a classifier trained on these data, random forest model fitting was performed and tuned until a satisfactory product was produced. The final trained classifier had a 7.6% out-of-bag error rate having sorted 41 of the 45 early-stage biofilms correctly and 44 of the 47 late-stage biofilms correctly (ntrees = 750, mtry = 31). This classifier was then used to sort the test dataset and its accuracy assessed. The classifier correctly sorted 29 of 31 test samples (14 of 15 early-stage and 15 of 16 late-stage biofilms; 93.5% accuracy).

Analyzing this classifier more closely, specific ASVs were assigned an importance value based on how they impact the accuracy of the classifier when they are removed (mean decrease in Gini). These importance values were then plotted for the top 25 most important ASVs and the taxonomic group they belong to was annotated (Supplementary Figure 6). Six of the top 25 most important ASVs belonged to the family Saprospiraceae (5 assigned to the family level and 1 belonging to the genus *Haliscomenobacter*), 4 belonged to the family Rhodobacteraceae (all assigned to the family level), 2 belonged to the family Verrucomicrobiaceae, and 2 belonged to the family Sphingomonadaceae (1 belonging to the genus *Sphingorhabdus* and 1 belonging to the genus *Rhizorhapis*). Additional genus-level ASVs in the top 25 included *Phreatobacter*, *Flavobacterium*, *Luteimonas*, *Peredibacter*, and the predatory myxobacterium *Nannocystis*. Of these, the *Haliscomenobacter* ASV had the highest genus-level importance value (mean decrease in Gini = 1.22).

To visualize these successional dynamics, relative abundances were plotted for taxa identified as important indicators by ANCOM-BC, random forest, or both, ordered from earliest to latest peak abundance (Figure 6). Early-stage biofilms were dominated by *Caulobacter* and *Flavobacterium*, both of which declined sharply after day 9. *Sphingorhabdus* peaked modestly at day 38 before declining. A distinct mid-succession cohort peaked at day 23: *Rhizorhapis* reached approximately 4.5% median relative abundance, *Bdellovibrio* peaked at approximately 0.4%, and *Methylotenera* reached approximately 3.2%, all declining thereafter. The Rhodobacteraceae rose rapidly from less than 1% at day 9 to approximately 6-9% by day 23 and remained elevated throughout the remaining time points. Late-stage biofilms were characterized by increasing Saprospiraceae (reaching approximately 18% by day 80), *Haliscomenobacter* (approximately 5% by day 80), Verrucomicrobiaceae (approximately 1%), and the predatory myxobacterium *Nannocystis*.

Fish pathogenic flavobacteria are of particular concern in cold water aquaculture facilities. At the species level, *Flavobacterium columnare* was detected at low relative abundance across all time points (35) (median < 0.1% at most time points) but was notably elevated at day 53 (median 0.35%, mean 0.73%, maximum 4.04%; Supplementary Figure 7). All 34 day 53 samples originated from raceways experiencing active columnaris disease (CD or CD/Gill Disease), and *F. columnare* abundance was highest in raceways R19 (mean 2.33%) and R18 (mean 0.60%) which had concurrent columnaris and gill disease. *Flavobacterium psychrophilum* was rarely detected (11 of 123 samples, median 0% at all time points; Supplementary Figure 8).

## Discussion

This study examined 19 rainbow trout aquaculture raceways at 6 varying time points over an initial 80-day hatchery production period where fish fry are grown up and then eventually transferred to outdoor raceway structures for grow out. The asynchronous nature of the raceway production period allowed our group to perform extensive sampling on a single trip and still capture a temporal picture of biofilm development on surfaces within these raceways. The picture we captured was one of discrete, distinct biofilm communities that change in composition over a matter of weeks in a predictable fashion.

All beta diversity metrics compared across the six time points sampled showed significant differences when looking at the factor of biofilm maturity overall as well as for all pairwise comparisons between the six time points. This is a strong signal that deterministic succession is occurring, with discrete time points mere weeks apart still yielding significantly different microbial community structures. When analyzing overall variance of these communities via beta dispersion analysis, early communities were shown to be significantly more variable than late-stage biofilms. This aligns with ecological successional theory, as early-stage founders appear and colonize somewhat stochastically depending on the individual raceway before converging towards a predictable mature biofilm community (48–52). Alpha diversity comparisons showed that the earliest sampled raceway, 9 days into its production period, had significantly fewer observed ASVs than raceways from 23 days onward, as well as significantly lower Shannon and Faith PD values. This points to two things, one being that colonization of surfaces by all members of a mature biofilm is not something that happens instantaneously; it requires both time and the likely creation of ecological niches within the biofilm structure that allow for colonization and proliferation. The second being that sometime between day 9 and day 23, the vast majority of the main mature biofilm members colonize and the overall community diversity stabilizes from this point through the final sampling point at day 80. This maturation of the biofilms occurs, despite efforts to manually remove the biofilms with brushes, suggesting that brushing is not completely effective for biofilm removal. However, it is important to note that while overall diversity may stabilize, the relative abundances of the various members continue to shift over each time point, as evidenced by the earlier observations regarding beta diversity.

Taxonomic resolution from a sequencing study can be limited by a number of factors including the size of the amplicon used, available reference sequences, and inherent taxonomic homogeneity for specific genes. This limitation is highlighted by some of the major taxonomic groups that comprise these biofilms, namely the families Saprospiraceae, Comamonadaceae, and Rhodobacteraceae. These families composed 25-50% of later stage biofilm samples. ASVs could not be assigned beyond the family level, again likely due to the aforementioned reasons. However, even without being able to assign these sequences to a genus or species, we are still able to differentiate between the individual ASVs. Every individual time point sampled had unique ASVs from these three major families. This strain dynamism is obscured when looking at the pure taxonomic data but becomes apparent when analyzing ASVs (Figure 4). This approach allowed us to observe not only that the plurality of ASVs are shared between five of the six time points (days 23 through 80), with nearly the same number shared across all six time points including day 9. This indicates that many taxa are not detected in the earliest biofilms but appear slightly later (by day 23).

A clear successional shift with specific bacterial groups representing early and late-stage biofilms was observed. Founder groups such as members of the Comamonadaceae as well as the genera *Flavobacterium* and *Rheinheimera* dominate the earliest stage biofilms sampled, comprising well over 50% of the community at day 9 (Figure 1B). These findings correlate well with other freshwater biofilm studies which also found *Flavobacterium* and *Rheinheimera* were primary colonizers that contributed to surface conditioning and niche building via EPS secretions and other mechanisms (14, 53). *Caulobacter* was another genus of bacteria flagged as early-enriched by both ANCOM-BC and random forest. This genus is a quintessential pioneer of freshwater biofilms (54). *Caulobacter* possesses a unique adhesin known as a holdfast, considered to be the strongest known biological adhesive, which would allow this genus to colonize a relatively clean, bare aquatic surface (55–58). *Pseudomonas* was also highly enriched in early-stage biofilms (59). This is a ubiquitous, r-strategist colonizer that consistently appears first in freshwater biofilm succession studies, secreting exopolysaccharides that will form the backbone of mature biofilms when *Pseudomonas* has long been outcompeted and replaced by late-stage biofilm members. *Sphingorhabdus* was also highly enriched in the early biofilm community. This genus of bacteria, along with the other members of the Sphingomonadaceae (*Sphingomonas*, *Sphingopyxis*, and *Blastomonas*) that were elevated, condition surfaces for subsequent colonizers through the production of glycosphingolipids.

The community then became more diverse by day 23 and the shifts in community dominance continued. Day 23 biofilms contained the highest percent abundance of *Bdellovibrio* and like organisms (BALOs) across all biofilm stages. BALOs are obligate predators of Gram-negative bacteria. At this early-mid biofilm state, a high percentage of the biofilm community is comprised of Gram-negative Proteobacteria pioneers. *Bdellovibrio* and *Peredibacter*, the BALOs found to be important indicators of biofilm state, decrease in abundance from this point onward. Previous studies have found that biofilm maturity provides a physical impediment to BALO predation, likely explaining the declining populations of BALOs as available prey becomes embedded in the EPS biofilm, becoming structurally protected (60).

A pair of interesting bacterial genera peak at day 23. *Rhizorhapis*, a member of the Sphingomonadaceae with nitrogen-fixing capacity, comprises close to 5% of the community at this early stage. This could potentially indicate nutrient input to the developing biofilm before being subsequently displaced as the biofilm matures. *Methylotenera* is an obligate methylotroph that uses methylamine as its sole energy source. Methylamine in this system likely derives from the demethylation of choline and glycine betaine released during lysis of pioneer bacteria, as membrane phospholipids and osmoprotectants are degraded by other community members (61–63). The peak abundance at day 23 of approximately 3% of the total community points to early colonizers beginning to die and initial nutrient cycles beginning.

In late-stage biofilms (day 53 and onward), the genus *Haliscomenobacter* returned the strongest enrichment signal across the entire dataset, being flagged in both ANCOM-BC analysis and as an important taxon in the random forest model we developed. *Haliscomenobacter* is a filamentous bacterium, previously described from active wastewater sludge (64). One can speculate that late-stage biofilms have accumulated a wide range of potential substrates as cells die and then degrade within the matrix. This created niche would then favor saprophytes such as *Haliscomenobacter*. This observation is also reinforced as the ASVs assigned to the family Saprospiraceae, comprised primarily of saprophytic microbes including *Haliscomenobacter*, became enriched in late-stage biofilms as well, increasing from nearly 0% of the community at day 9 to around 18% of the community by day 80. *Nitrosomonas* was also a late-enriched microbe with ammonia-oxidizing capabilities. *Nitrosomonas* are slow-growing and require sufficient ammonium concentrations, thus likely explaining their late increases. Ammoniaoxidizing bacteria are important in the context of a fish hatchery as the nitrification process can directly combat ammonia toxicity which is a threat to fish stock. This suggests that mature hatchery biofilms may confer a functional benefit to water quality by contributing to biological nitrification. Notably, this successional progression toward a community enriched in ammonia-oxidizing and saprophytic taxa occurred despite routine scrubbing of raceway walls to remove visible biofilm, indicating that established biofilm communities are resilient to mechanical disturbance. These observations suggest that biofilm management practices should consider not only the potential risks of mature biofilms (e.g., pathogen harboring) but also their potential benefits for water quality.

In addition to saprophytic microbes, an interesting group of predatory bacteria became enriched in late-stage biofilms. *Nannocystis*, *Polyangium*, and *Haliangium*, all belonging to the predatory myxobacteria, were flagged as differentially abundant in mature biofilms. Myxobacteria engage in a so-called “wolfpack” behavior where large swarms of these microbes secrete lytic enzymes cooperatively (65). High prey densities and complex biofilm architectures favor these specific microbes.

Overall, we can see unique and interesting ecological succession occurring in the biofilms within this fish hatchery environment. Early founder species like *Pseudomonas*, *Caulobacter*, and *Flavobacterium* arrive, modify the surface environment, fade, and establish habitat and metabolic niches for late-stage species like the Saprospiraceae including *Haliscomenobacter*. Notably, these early-stage pioneer taxa (*Pseudomonas*, *Flavobacterium*, and other Gammaproteobacteria) correspond closely to the organisms most readily recovered by culture-dependent methods from hatchery surfaces in a previous study of this facility (66), suggesting that standard culturing approaches may systematically underrepresent the diversity of mature biofilm communities. Young biofilms are enriched in solitary, predatory BALOs whereas mature biofilms are enriched with social, predatory myxobacteria. Nitrogen cycling indicators like *Rhizorhapis* (nitrogen fixation), *Methylotenera* (denitrification), and *Nitrosomonas* (nitrification) rise and fall depending on substrate availability within the biofilm microenvironment.

An important consideration when interpreting amplicon sequencing data is that relative abundance changes do not necessarily reflect changes in absolute abundance. However, qPCR estimates of total bacterial load demonstrated a significant positive correlation with biofilm age (Spearman ρ = 0.50, p < 0.001), with late-stage biofilms harboring approximately 2.5-fold more bacteria than early-stage biofilms. This confirms that the biofilm community genuinely expands over time and that late-enriched taxa such as *Haliscomenobacter* and Saprospiraceae represent real increases in both relative and absolute terms. Conversely, the decline of early colonizers such as *Caulobacter* and *Pseudomonas* likely reflects genuine displacement rather than simple dilution, as these taxa decreased in relative abundance far more than the approximately 2.5-fold increase in total biomass could explain. Furthermore, ANCOM-BC employs a log-ratio framework with bias correction specifically designed to address compositionality in amplicon data, lending additional confidence to the differential abundance results reported here.

Several limitations of this study should be acknowledged. First, the experimental design is cross-sectional rather than longitudinal: biofilms of different ages were sampled from different raceways rather than tracking individual raceways over time. While this approach captures raceway-to-raceway variability and provides large sample sizes per time point, raceway-specific effects cannot be entirely excluded. Indeed, within-timepoint PERMANOVA testing confirmed that raceway identity contributed significant compositional variation at most time points (p < 0.05 for five of six time points; data not shown), particularly at days 38 and 53 where differing disease histories among raceways likely contributed to this effect. Nevertheless, all pairwise maturity comparisons remained highly significant (p < 0.001), indicating that the successional signal is robust to raceway-level variability. Second, late-stage raceways had undergone chemical treatments (salt, hydrogen peroxide, and/or chloramine-T) during the production period, and some community shifts at later time points may partly reflect treatment history rather than succession alone. Relatedly, because all sampled raceways were actively stocked, biofilm age is confounded with cumulative nutrient and organic loading: feed input and fish biomass increased over the production period, so older biofilms necessarily developed under progressively higher nutrient availability. The positive correlation between total bacterial load and biofilm age reported above is consistent with this. Our observational design therefore cannot fully distinguish autogenic succession, driven by the biofilm community itself, from nutrient-driven (allogenic) change; cleanly separating the two would require sampling biofilms developed in raceways supplied with hatchery water but without fish, which was not feasible in this working facility. Third, the V4 region of the 16S rRNA gene does not resolve many taxa beyond the family level, particularly within the Saprospiraceae, Rhodobacteraceae, and Comamonadaceae that comprise substantial portions of these biofilms. The ASV-level strain dynamism observed in Figure 4 likely underestimates true strain turnover, and shotgun metagenomics would be needed for finer resolution. Additionally, 16S rRNA gene copy number varies among bacterial taxa, which means that qPCR-based estimates of total bacterial load are inherently weighted toward high-copy-number organisms and do not directly reflect cell counts. The swab-to-discoloration sampling protocol also means that the surface area sampled was not standardized, so absolute inter-sample comparisons of bacterial load should be interpreted as estimates rather than precise measurements. Finally, the rarefaction depth of 8,000 reads, while necessary to retain samples with lower sequencing depth, may have excluded rare taxa that could affect the unique ASV counts at less deeply sampled time points.

*Flavobacterium* species are among the most consequential bacterial pathogens in cold-water aquaculture, with *F. columnare* causing columnaris disease and *F. psychrophilum* causing bacterial cold water disease, together responsible for significant morbidity and economic losses in salmonid hatcheries worldwide (67). The genus *Flavobacterium* was the single most abundant taxon in day 9 biofilms (median 34% relative abundance) and was significantly enriched in early-stage communities by ANCOM-BC (LFC = -1.04). At the species level, *F. columnare* was detected at low abundance throughout the production period but was notably elevated at day 53, where all sampled raceways were experiencing active columnaris disease outbreaks (35). This confound prevents disentangling whether the elevated *F. columnare* signal reflects biofilm succession or pathogen shed from infected fish colonizing the biofilm. Nevertheless, the persistent low-level detection of *F. columnare* across biofilm stages suggests that hatchery biofilms may serve as a reservoir for this pathogen, consistent with previous reports of *F. columnare* persistence in aquatic biofilms (68). *F. psychrophilum*, by contrast, was essentially absent from biofilm communities (detected in only 11 of 123 samples), suggesting that biofilms in this system do not serve as a meaningful reservoir for cold water disease, consistent with earlier findings from this facility (34).

The incomplete effectiveness of mechanical cleaning has practical implications for hatchery management. Routine brushing reduces biofilm biomass but, as shown here, does not prevent rapid re-colonization or the re-establishment of a characteristic mature community within roughly two to three weeks, indicating that brushing resets standing biomass without altering the successional endpoint. Where biofilms can act as reservoirs for fish pathogens such as *F. columnare*, this suggests that brushing alone is unlikely to be sufficient for biosecurity and may need to be combined with disinfection or more frequent intervention. This study was not designed to test cleaning efficacy directly; controlled before-and-after or brushed-versus-unbrushed comparisons would be needed to evaluate it rigorously.

Taken together, these findings demonstrate that biofilm communities on hatchery raceway surfaces undergo rapid, predictable ecological succession characterized by deterministic turnover from r-strategist pioneers to a diverse, mature community enriched in saprophytes, predatory myxobacteria, and nitrifying bacteria. The predictability of this succession, as evidenced by the high accuracy of our random forest classifier, suggests that biofilm age could serve as a practical indicator of raceway condition. In a management context, significant deviations from the expected successional trajectory, such as persistent dominance of pioneer taxa in an ostensibly mature biofilm, or unusual enrichment of pathogen-associated genera, could signal disrupted biofilm development or emerging health concerns warranting investigation. Moreover, the detection of fish pathogens such as *F. columnare* within biofilm communities, even at low abundances, underscores the importance of understanding biofilm ecology for disease management in aquaculture. Future work incorporating shotgun metagenomics and longitudinal sampling of individual raceways would provide finer taxonomic resolution and allow direct tracking of community trajectories, while functional characterization of biofilm communities could reveal whether the successional patterns described here have direct consequences for fish health and water quality.

## Data and Code Availability

Raw sequence data have been deposited in the NCBI Sequence Read Archive (SRA) under BioProject accession PRJNA1477683. Analysis code is available at https://github.com/todd-testerman/hatchery-biofilm-succession.

## Acknowledgments

We would like to acknowledge Brandon O’Sullivan, Alejandra Jáquez and Angeline Casale for helpful comments on the manuscript. We would also like to acknowledge Molly Schiffer and Colleen Brown for their work on DNA extractions.

