## Supplementary Figure for "Succession and Shifting Identities in Freshwater, Built Environment Biofilm Communities"

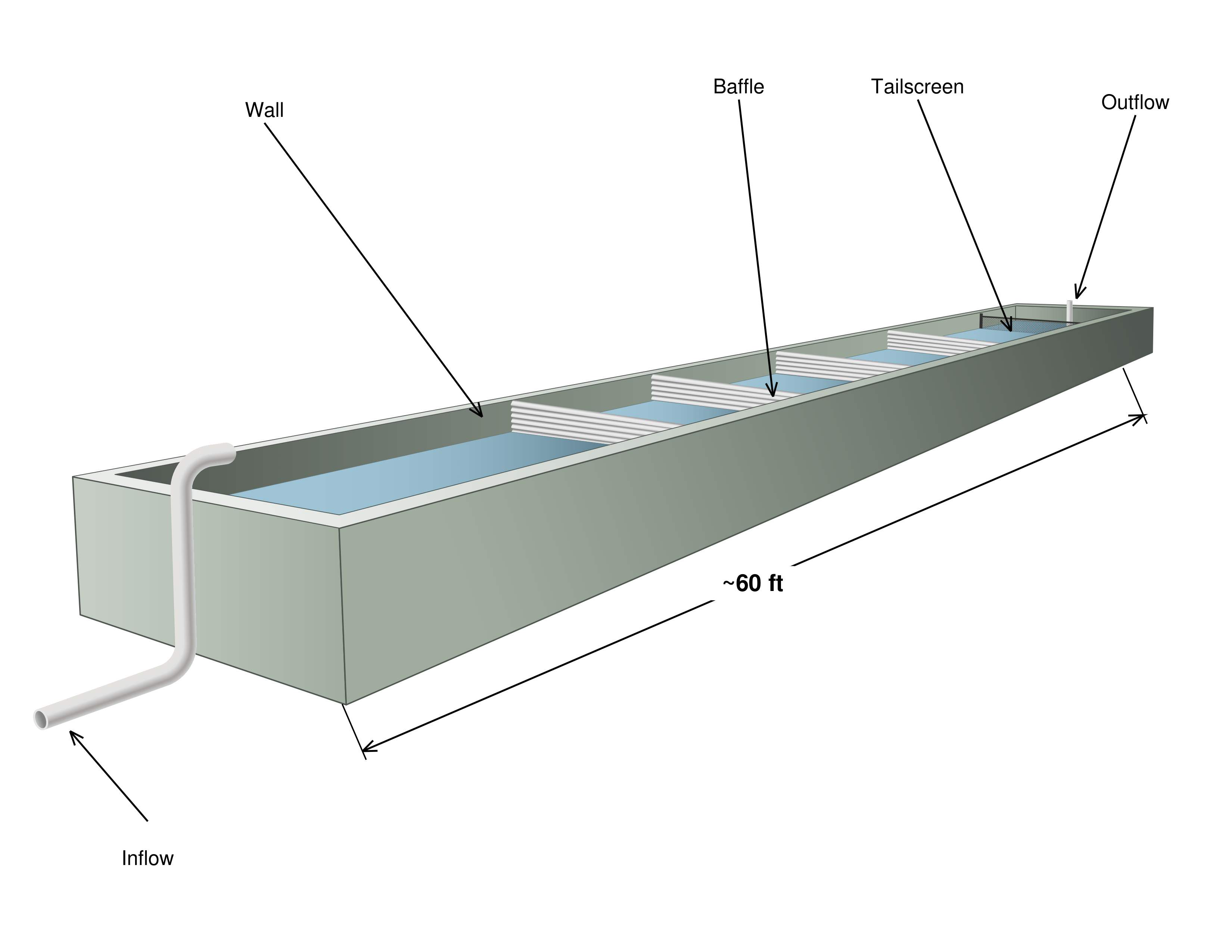

**Supplementary Figure 1**: Hatchery Raceway Diagram. Three-dimensional schematic of a typical epoxy-coated concrete hatchery raceway showing the sampled surfaces (walls, baffles, and tailscreen) along with the inflow and outflow positions. The scale bar indicates the approximate raceway length (~60 ft).

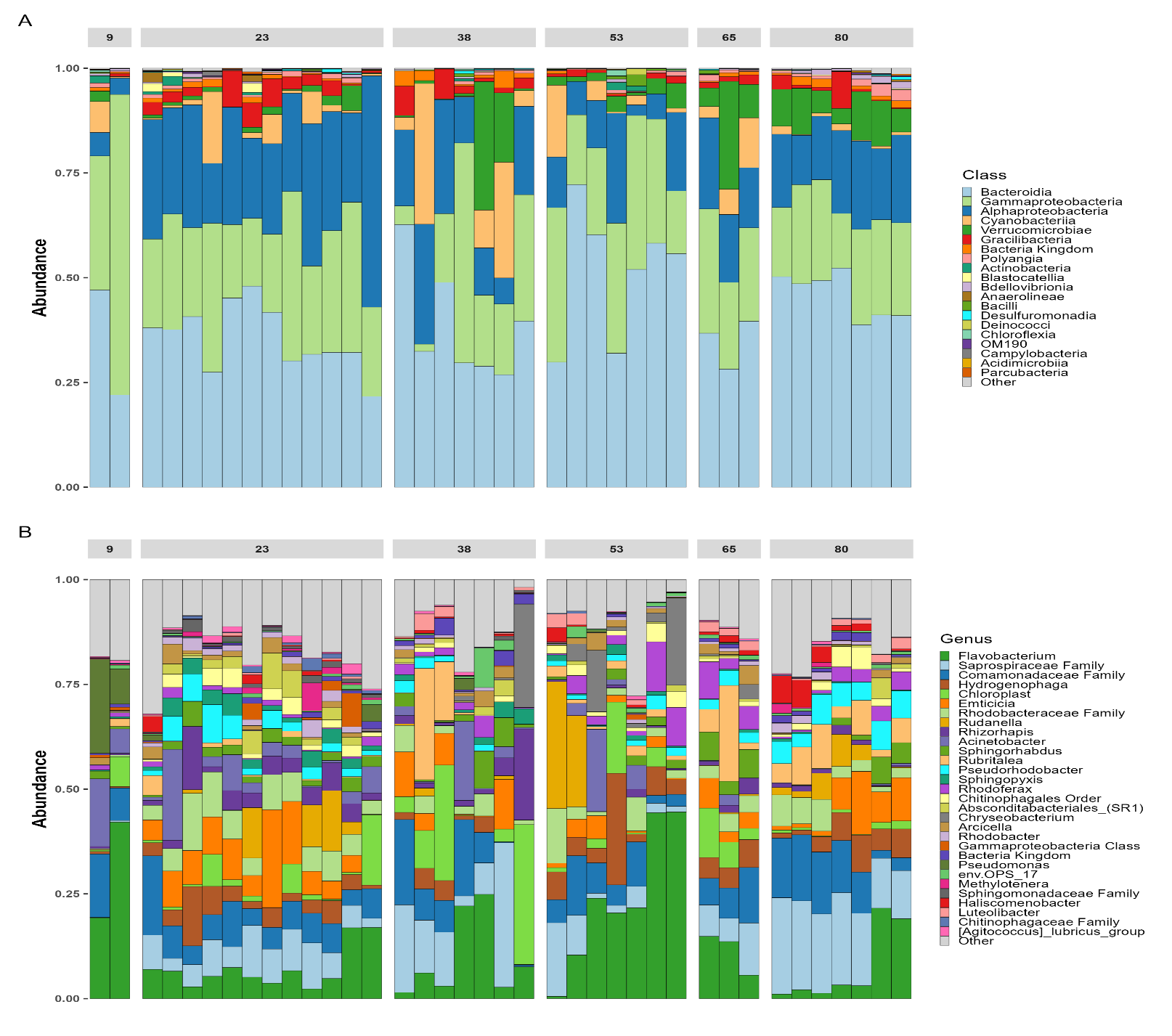

**Supplementary Figure 2: Taxonomic Composition of Tailscreen Biofilm Communities.** (A) Class-level and (B) genus-level relative abundance profiles of biofilm communities collected from tailscreen surfaces, faceted by biofilm age (days). The top 20 classes and top 30 genera are shown; remaining taxa are grouped as "Other." Samples within each timepoint are ordered by Bray-Curtis dissimilarity. Color palettes match those used in Figure 1 for direct comparison with wall swab communities.

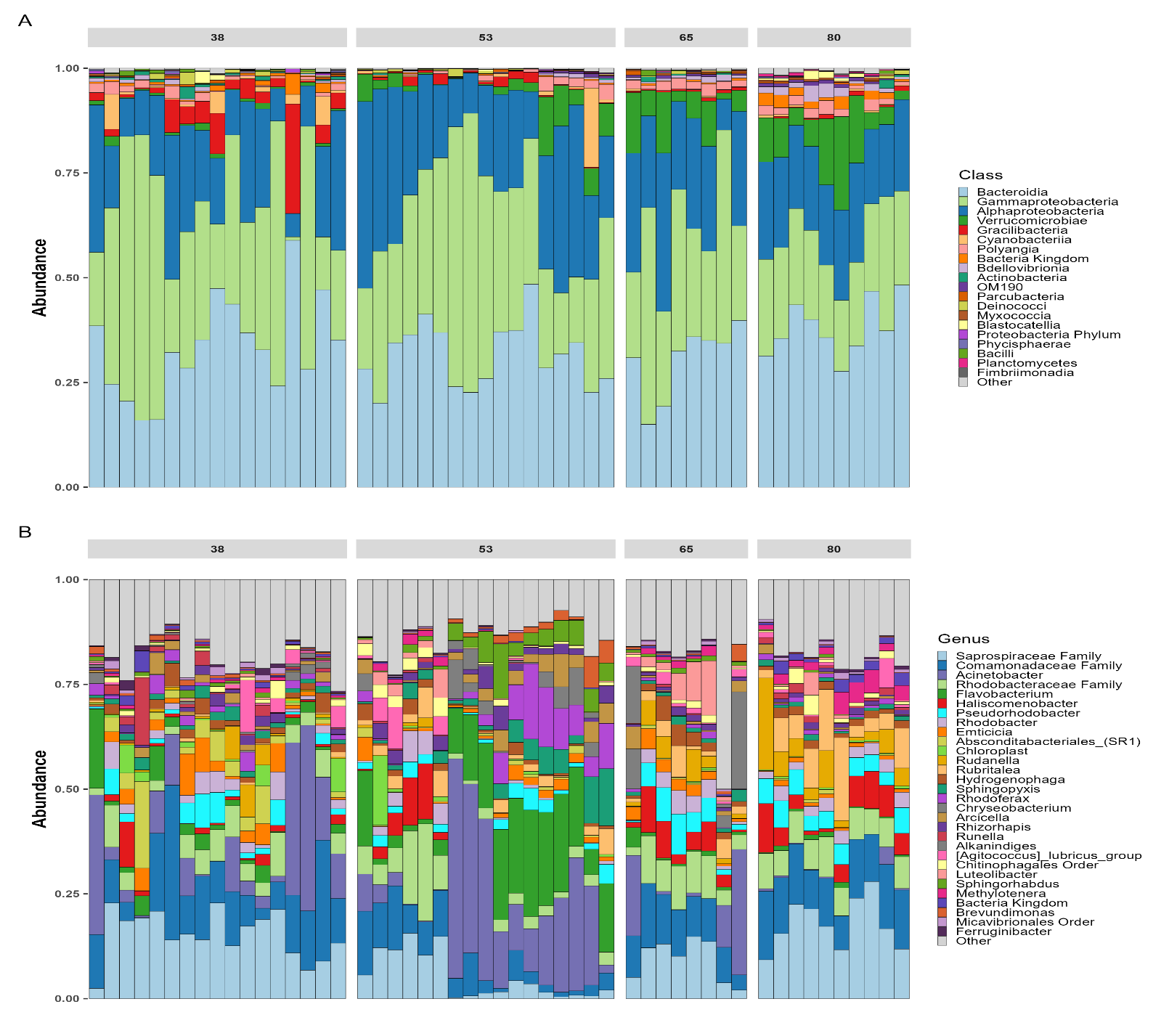

**Supplementary Figure 3: Taxonomic Composition of Baffle Biofilm Communities.** (A) Class-level and (B) genus-level relative abundance profiles of biofilm communities collected from baffle surfaces, faceted by biofilm age (days). The top 20 classes and top 30 genera are shown; remaining taxa are grouped as "Other." Samples within each timepoint are ordered by Bray-Curtis dissimilarity. Color palettes match those used in Figure 1 for direct comparison with wall swab communities.

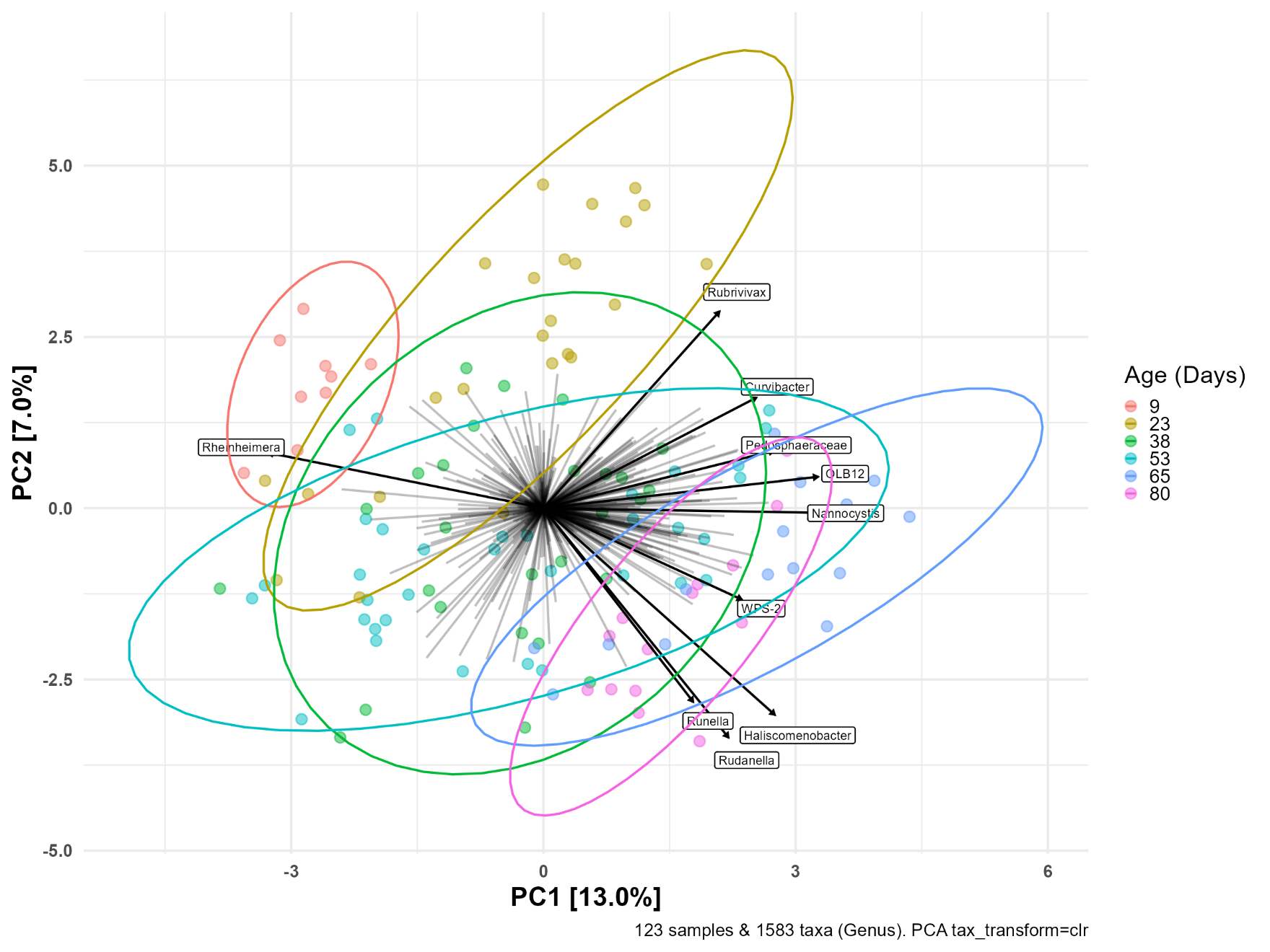

**Supplementary Figure 4: PCA Biplot with Top 10 Taxonomic Drivers.** Principal component analysis of wall swab biofilm communities using centered log-ratio (CLR) transformed genus-level abundances. Arrows indicate the direction and magnitude of the 10 genera contributing most to ordination structure. Points are colored by biofilm age (days), with 95% confidence ellipses shown for each timepoint. Unrarefied data were used for CLR transformation.

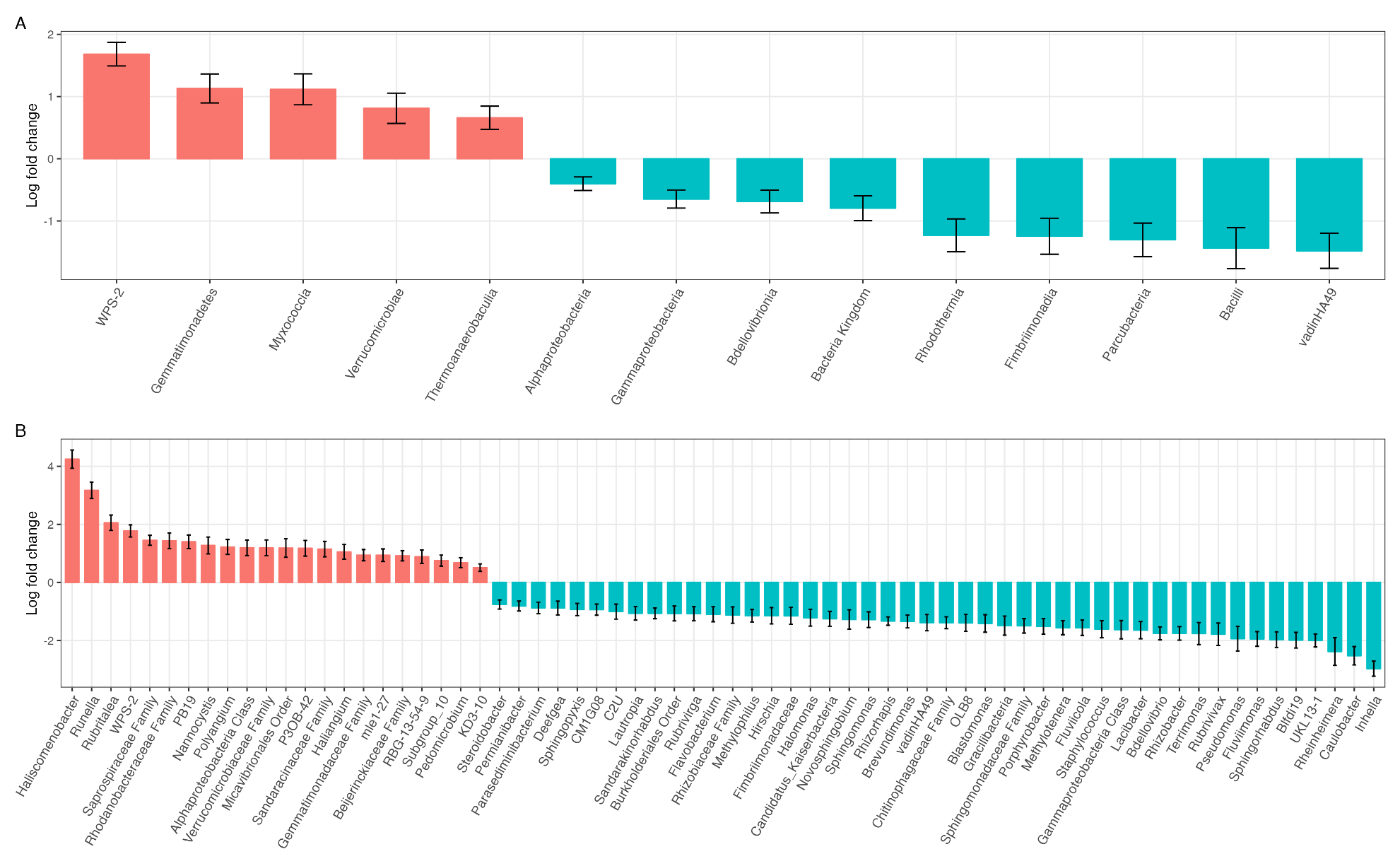

**Supplementary Figure 5: Class-Level UpSet Plot of Shared and Unique ASVs.** UpSet plot displaying the number of ASVs shared between and unique to each biofilm age group, with bars colored by class-level taxonomy (top 10 classes shown). This complements the genus-level UpSet plot in Figure 4. Set sizes (left) indicate the total number of ASVs detected at each timepoint. Data were merged by timepoint and rarefied to 8,000 reads per sample prior to merging.

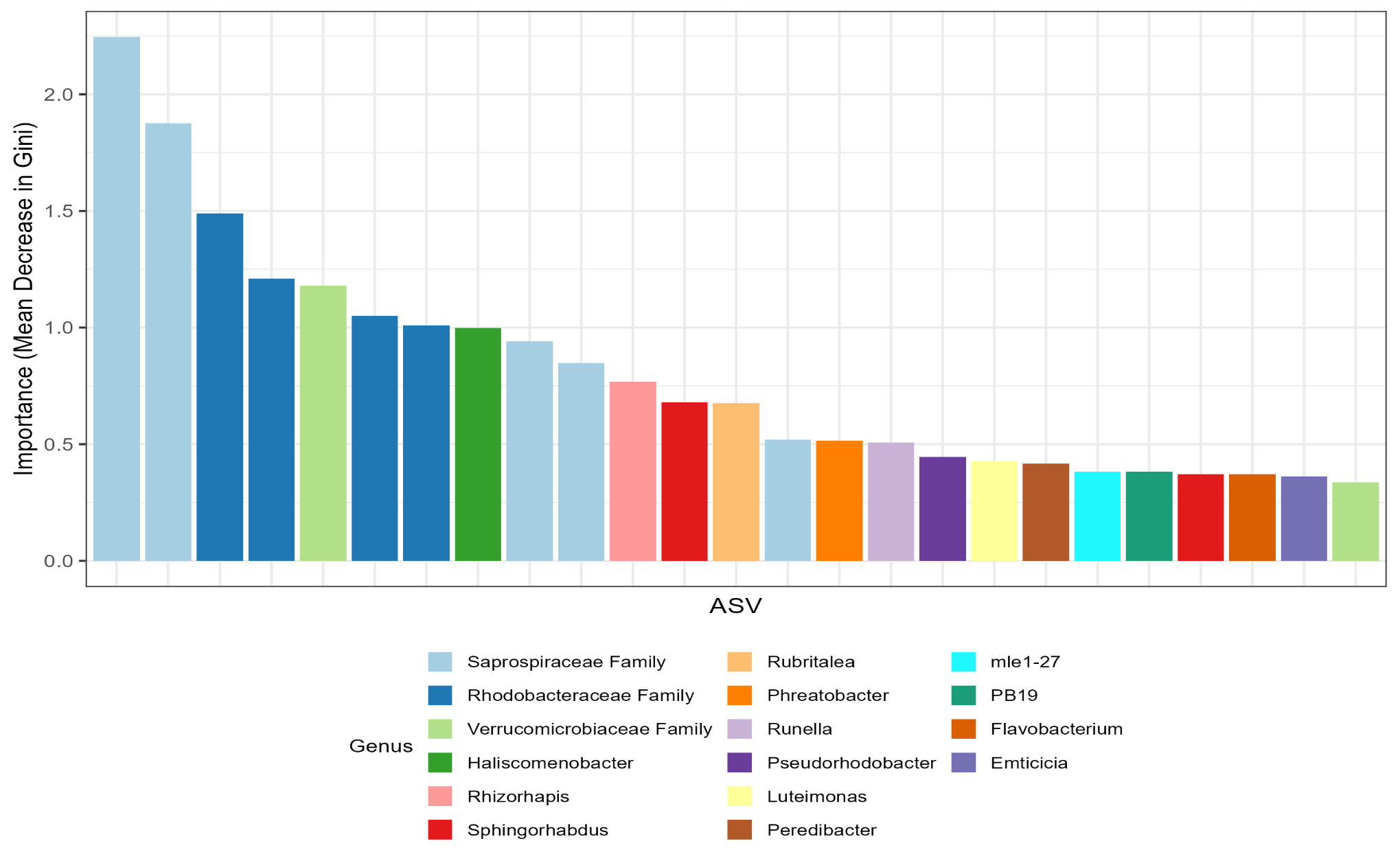

**Supplementary Figure 6: Top 25 ASVs by Random Forest Importance.** Mean Decrease in Gini importance scores for the 25 most informative ASVs in the random forest classifier distinguishing early-stage (days 9-38) from late-stage (days 53-80) wall swab biofilm communities. Bars are colored by genus-level taxonomy. The model was trained with 750 trees on an 80/20 stratified split of samples filtered to ASVs present in at least 10% of samples.

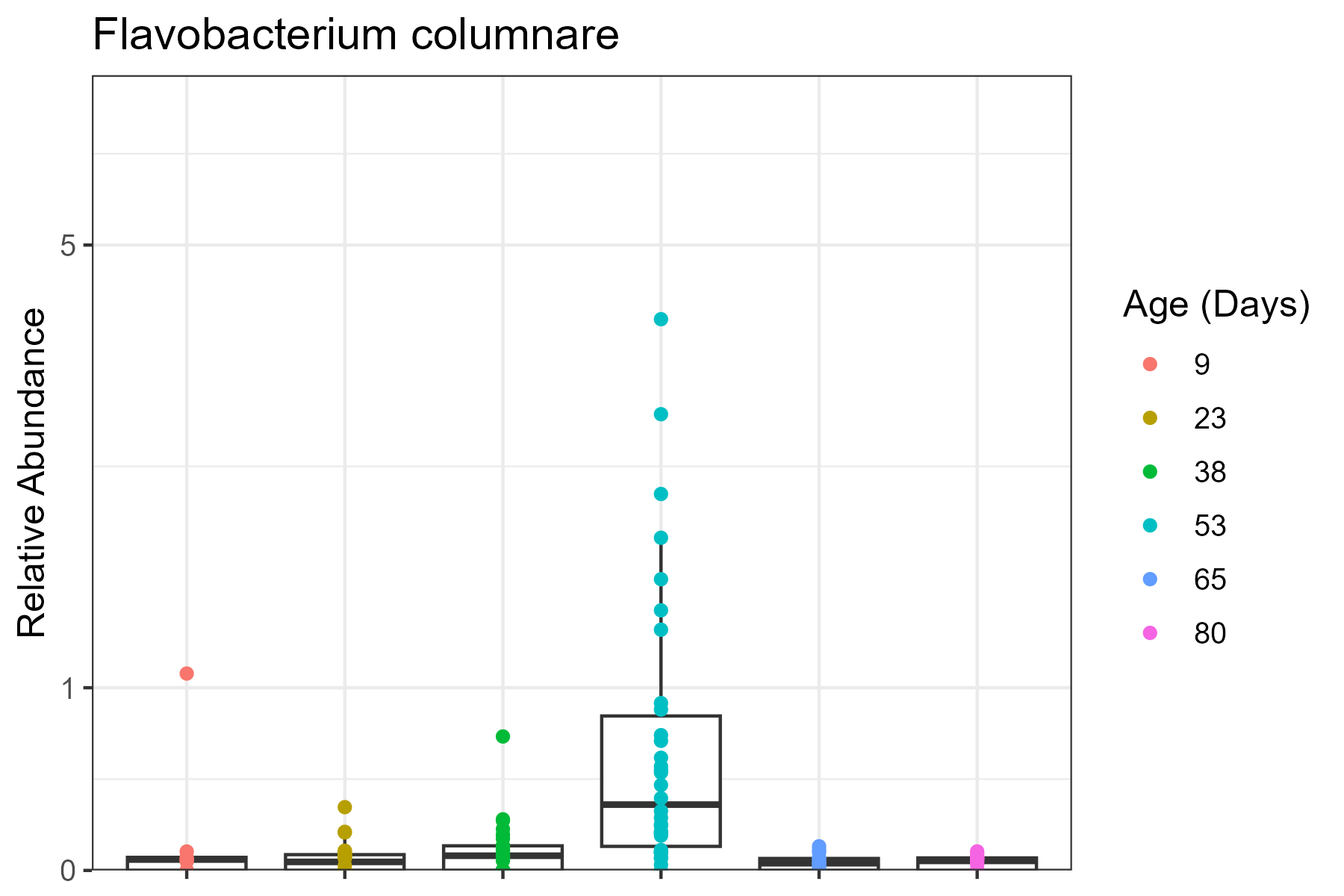

**Supplementary Figure 7: Relative Abundance of *Flavobacterium columnare* Across Biofilm Age.** Species-level relative abundance (%) of F. columnare in wall swab biofilm communities at each sampling timepoint. *F. columnare* was detected at low levels throughout the study but showed elevated abundance at day 53, when all sampled raceways had active columnaris disease outbreaks. Y-axis uses pseudo-log scaling.

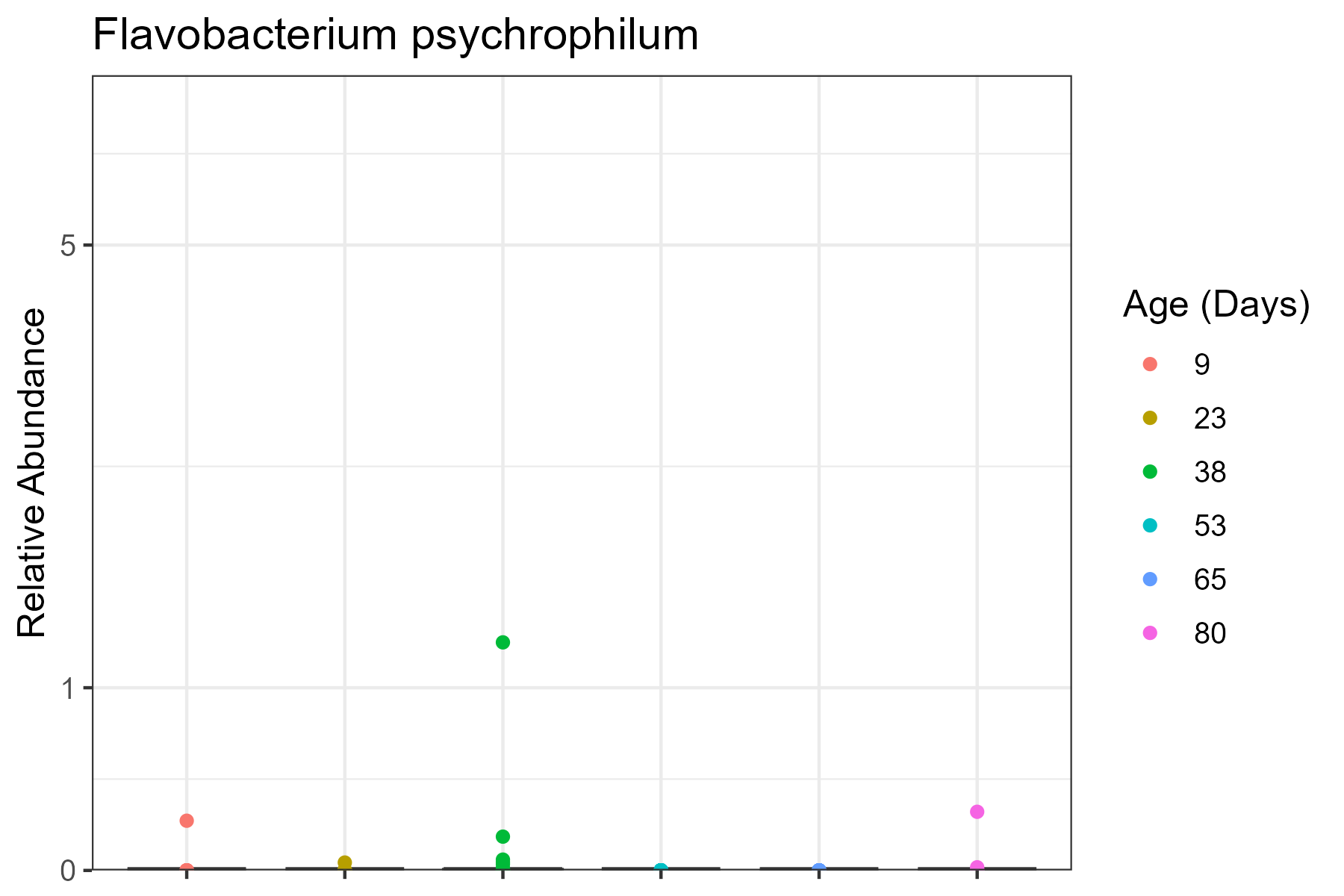

**Supplementary Figure 8: Relative Abundance of *Flavobacterium psychrophilum* Across Biofilm Age.** Species-level relative abundance (%) of F. psychrophilum in wall swab biofilm communities at each sampling timepoint. *F. psychrophilum* was detected in only 11 of 123 samples and was essentially absent from biofilm communities throughout the study period. Y-axis uses pseudo-log scaling.
